# Antiviral effect of thiazolides relies on mitochondrial mild uncoupling

**DOI:** 10.1101/2022.09.16.508272

**Authors:** Noureddine Hammad, Céline Ransy, Benoit Pinson, Jeremy Talmasson, Christian Bréchot, Frédéric Bouillaud, Jean-François Rossignol

**Affiliations:** Institut Cochin, INSERM, CNRS, Université de Paris, F75014 Paris, France; Service analyses Métaboliques-TBMcore, Université Bordeaux - CNRS UAR 3427 - INSERM US005, Bordeaux, France; Romark Institute of Medical Research, Tampa, FL, United States; University of South Florida, College of Medicine, Tampa, FL, United States; Global Virus Network, United States

## Abstract

**Background:** Viruses are dependent on cellular energy metabolism for their replication, the drug Nitazoxanide (Alinia) was shown to interfere with both. An effect of Alinia on cellular energy metabolism is the uncoupling of mitochondrial oxidative phosphorylation (OXPHOS). Our hypothesis was that uncoupling grounds the antiviral properties of Alinia.

**Methods:** Alinia or an unrelated uncoupler were applied to a viral releasing cell line to obtain the same increasing levels of uncoupling hence identical interference with OXPHOS.

**Findings:** Decrease in infectious viral particles release reflected the intensity of interference irrespective of the nature of the drug and was significant with modest deviation (≤25%) from normal.

**Interpretations:** A mild interference on cellular energy metabolism impacts significantly on viral replication cycle. This would explain Alinia’s antiviral properties *in vitro* moreover antiviral action of Alinia is supported by clinical trials.

**Perspectives:** Altogether this indicates that moderate interference with mitochondrial bioenergetics should be considered as a ground for a therapeutic effect. In addition, Alinia would constitute example for a safe therapeutical use of an uncoupler, which deserves consideration for a wider range of applications.

## Introduction

### Nitazoxanide interferes with cellular energy metabolism

Nitazoxanide (NTZ) is a synthetic thiazolide developed in the 90’s against new opportunistic protozoan infections in AIDS patients (Fig. 1). In aqueous solution (as plasma), NTZ is within 15-30 min deacetylated into tizoxanide (TZ), the active metabolite. Tizoxanide (TZ) is a weakly polar molecule and high protein-binding, more than 99.9% of circulating TZ is bound to the plasma proteins. Reported concentrations of TZ in plasma of treated patients range from a peak value of 2mg/L (7.5µM) after single dose of 500mg^1^ to values ranging between 10 and 100µM under repetitive high dosage^2^. In the last 20 years, NTZ was subject to many screening studies with aim to explore new therapeutic applications. Evidences raised the issue of interference of Nitazoxanide with cellular energy metabolism and consequently a legitimate question is the nature of the relations between these observations and the therapeutic properties of NTZ.

**Figure 1:**
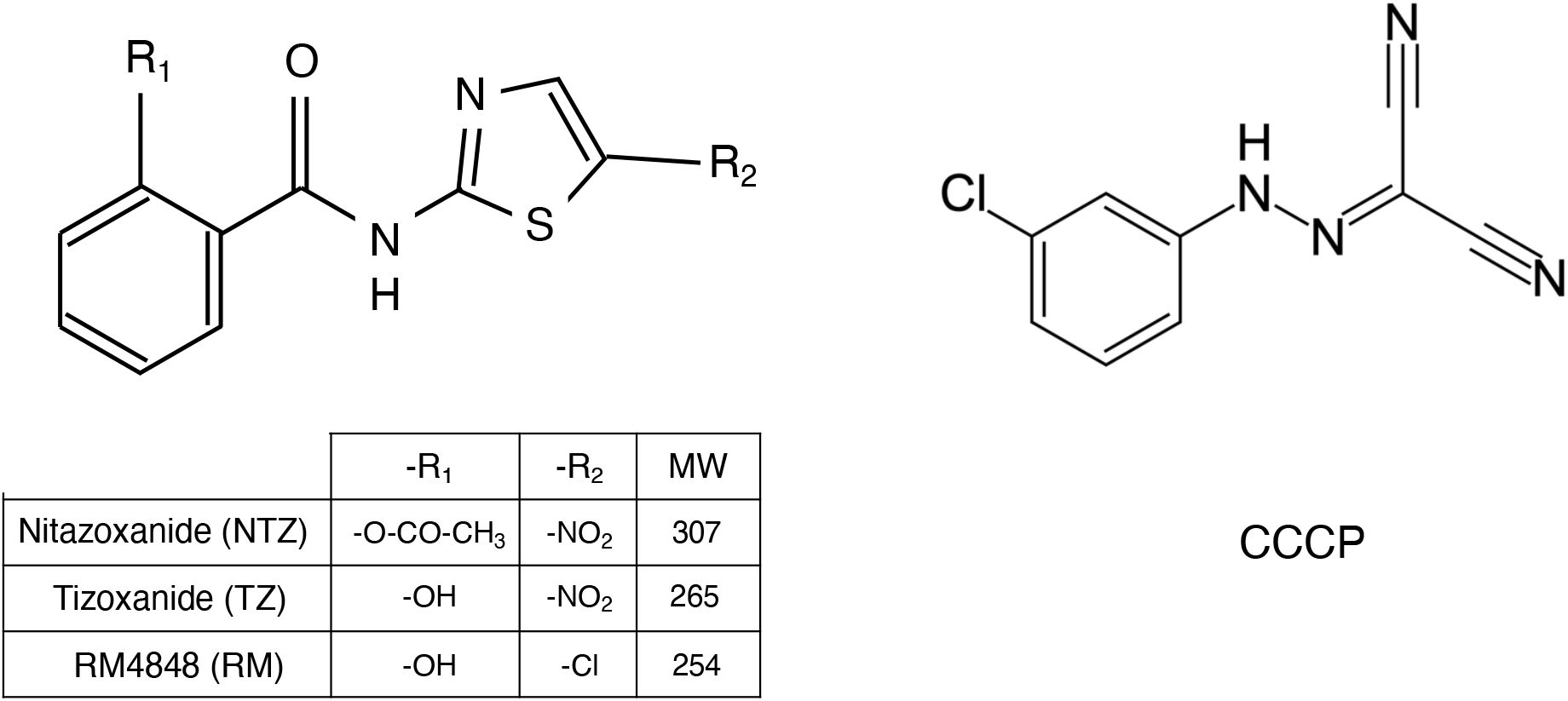
Nitazoxanide, Tizoxanide, RM and the uncoupler Carbonyl cyanide m-chlorophenyl hydrazone (CCCP).

A first proposal was that the therapeutic effect of NTZ relies on the specific properties of a metabolic enzyme found in the target species: the pyruvate ferredoxin oxidoreductase (PFOR)^3^. This enzyme is essential for their anaerobic energy metabolism and is not found in mammals. This effect would rely on the similarity between NTZ and Thiamine pyrophosphate (TPP) a cofactor used by PFOR. This TPP is also a cofactor for mammalian pyruvate dehydrogenase (PDH) and inhibition of mammalian PDH was not examined. Another issue is the presence of the nitro group that was apparently required for inhibition of PFOR while derivatives without this nitro group showed similar anti parasitic activities^4^. Therefore, indication for absence of a metabolic interference of the same nature in mammalian cells is lacking and effect through PFOR inhibition in target species remains uncertain.

The second type of interference proposed for NTZ refers to the uncoupling of cellular respiration. The main role for cellular respiration is the oxidative phosphorylation (OXPHOS) that regenerates continuously the ATP whose hydrolysis feeds the cellular energy requirements. in most animals OXPHOS is the largest contributor to cellular energy metabolism. Beside OXPHOS part of ATP is generated by substrate linked phosphorylation steps during glycolysis or TCA cycle. In contrast with these enzymatic reactions, OXPHOS is not stoichiometric and therefore the actual number of ATP generated differs from the theoretical prediction and depends on the yield of conversion between substrate oxidation and ATP formation. Two types of interferences might therefore be expected: either the OXPHOS is impaired by poisoning or the yield of conversion is deteriorated and this is how uncouplers act. The molecular explanation is that uncouplers increase the passive proton conductance of the mitochondrial inner membrane (chemiosmotic theory of Peter Mitchell). The cell/animal aims to compensate the deterioration of yield by an increased respiratory activity and, when possible, it would simply result in an increase in energy expenditure. If this is not possible toxic effects are expected. Actually, the first uncoupler (dinitrophenol) was used to promote weight loss^5^. Severe adverse effects appeared and for this reason dinitrophenol was banished from use^6^. Consequently, uncoupling is rather considered as an unwanted side effect for a drug. At this step it should be highlighted that the safety record of NTZ is excellent with little unwanted side effects.

Studies in M. tuberculosis have shown that NTZ acts as an uncoupler disrupting membrane potential and intra-organism pH homeostasis^7^. A High-throughput screening of FDA-approved drugs study^8^ demonstrated that NTZ increased oxygen consumption of cells. Consequently, it was demonstrated that NTZ prevents MPP^+^-induced block in mitochondrial respiration and restores ATP production and while this seems rather counterintuitive, this was attributed to its mild uncoupling effect on oxidative phosphorylation^9^. NTZ has been considered for Alzheimer disease^10^ because TZ promotes autophagy process in BV2 cells, SH-SY5Y cells and primary neuron cells and primary astrocytes. through downregulation of PI3K/AKT/mTOR and NQO1/mTOR pathways^11^ NTZ downregulate the activity of GSK3β trough downregulation of PI3K/AKT. Clinical trials evidenced that NTZ improved the outcome of viral infections with influenza^12^, hepatitis B^13^ viruses and preliminary results were obtained for COVID-19^14 15^. In vitro studies demonstrated a wide range of viruses whose infectious cycle was affected by NTZ: Hepatitis B and C viruses^16^, influenza^17^, HIV^18^ and Sars-Cov-2^19^. This suggests that NTZ action targets the eucaryotic host cell with little influence of the nature of the virus.

### Hypothesis: partial uncoupling explains the antiviral effect of nitazoxanide

Our hypothesis was therefore that the antiviral action of NTZ relies on its uncoupling activity. It leads immediately to two predictions *1)* if the uncoupling effect of a reference mitochondrial uncoupler and of TZ are matched the impact on the viral replication process would be identical for both drugs. In other words: the intensity of interference with mitochondrial respiration would be the relevant issue and not the nature of the drug. *2)* This mechanism is expected to be of general importance and should be observable also with non-pathogenic viruses such as those commonly used as viral gene expression vectors produced by engineered packaging cell lines. In addition to these predictions, a requirement was that the deleterious effect against virus should not result from death of the virus replicating cells and therefore should take place with mild levels of interference with mitochondrial function. With regard to the scenario explaining how uncoupling would impair virus replication before cell death, the most likely relies on the facts that on one side all steps of virus replication are dependent on energy, and on the other side, cells subject to energy shortage trigger a re-orientation of the energy use towards priorities^20 21^ and viral replication would be adversely affected by this re-orientation.

## Results

### Inhibition of cellular oxidative phosphorylation with thiazolides and the reference uncoupler CCCP

The cell line used hereafter was the Phoenix ECO cell line an ecotropic retroviral packaging cell line susceptible to release infective (but non-replicating) retroviral particles after transfection by a DNA sequence for a retroviral expression vector. In presence of glucose, the cellular ATP turnover is ensured both by respiration and by lactic fermentation. The suppression of ATP synthase activity by oligomycin is a widely used strategy to evaluate relative contribution of OXPHOS to cellular ATP turnover. In these cells the non-mitochondrial contribution to cellular oxygen consumption was negligible (not shown) and therefore the oxygen consumption rate (OCR) observed after oligomycin quantifies the mitochondrial respiration not coupled to ATP generation (proton leak). Addition of uncouplers (TZ, RM, or CCCP) increased the cellular OCR in presence/absence of oligomycin (Fig. 2A-C) up to a maximal respiratory rate that could not be increased further and with a same maximal value in presence/absence of oligomycin. The difference between the OCR in presence/absence of oligomycin (the grey area in Fig. 2A-C) represented an OCR explained by the activity of the ATP synthase and therefore provided a quantitative estimation for the OXPHOS rate. It decreased gradually to reach zero when OCR was maximal, with further uncoupler additions the OCR remained stable or declined and then the difference in presence/absence of oligomycin should refer to other explanations than oxygen consumption by OXPHOS. The impact of uncouplers on OXPHOS was therefore considered maximal (100% of interference with OXPHOS) when the dosage had the consequence that no effect of oligomycin on OCR could be detected anymore. This tallies with a simple model in which the OCR rate cannot be increased indefinitely because of enzymatic limitations with the consequence that leak and OXPHOS are competitors for use of this OCR. Then estimation of OXPHOS rate could be represented in relative units according to drug concentration (Fig. 2D-F).

**Figure 2:**
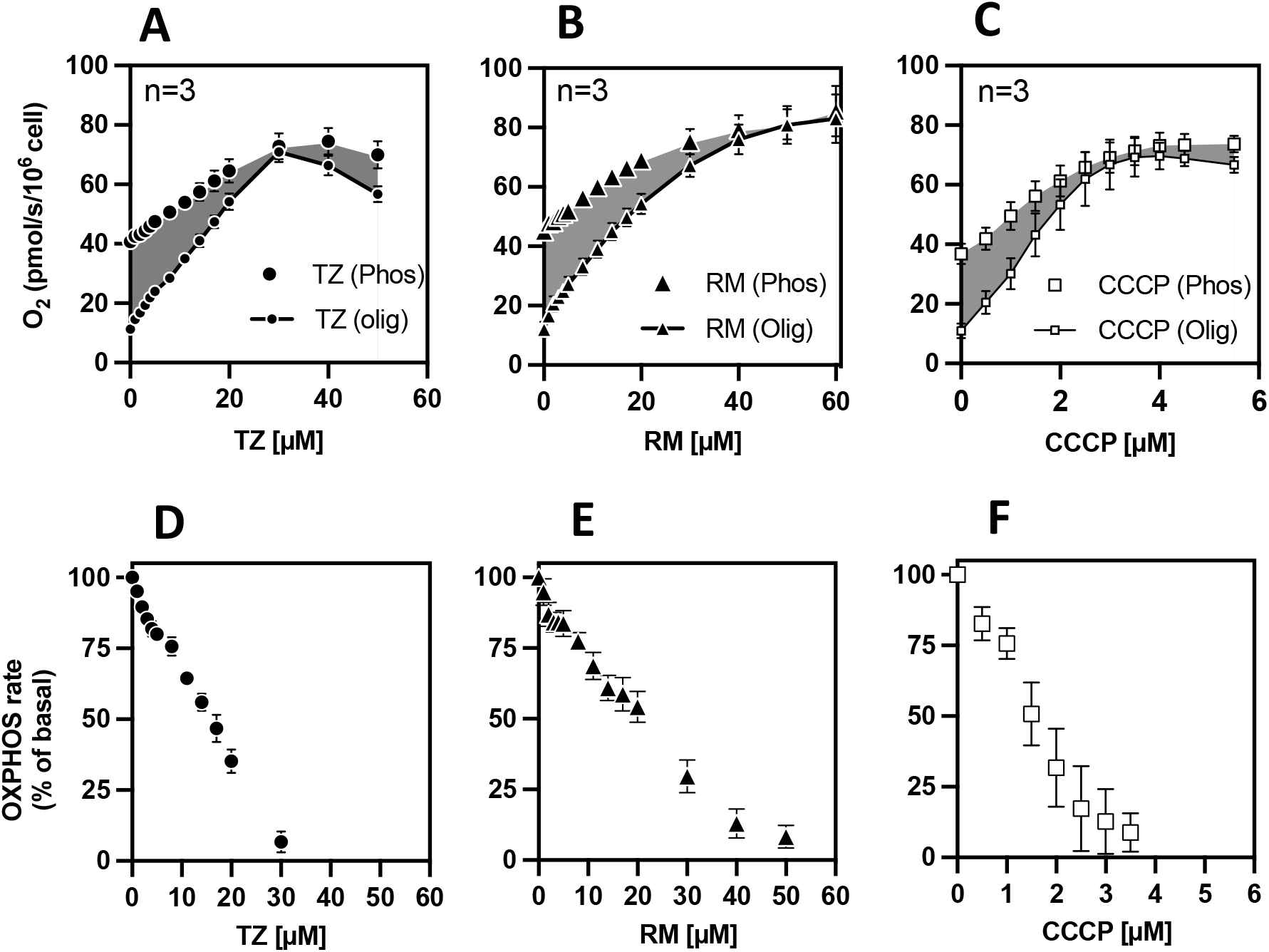
TZ, RM or CCCP effect on the cellular OCR of Phoenix ECO cells. Data in A, B and C are expressed as mean (X) in picomole of oxygen molecules per second and per 10^6^ of cell ± SEM) of minimum three independent experiments. Filled symbols and empty symbols are data of OCR in phosphorylating state and non-phosphorylating state respectively. The OCRs due to the activity of ATP synthase in D, E and F were calculated as described in Materials and Methods.

This led to determination of concentrations of TZ, RM or CCCP that causes stationary states with 25, 50, 75, and 100% of interference with OXPHOS that were used in subsequent experiments. A conclusion from this study is that the two thiazolides were approximately ten times less efficient than CCCP on this cellular model (X axis of Fig. 2).

### Glycolytic compensation to defective oxidative phosphorylation

Cell survival and proliferation are good indicators of the energy homeostasis of the cell, cell viability assays would evaluate both simultaneously. To investigate the ability of the Phoenix ECO cell line to use glycolysis to compensate OXPHOS impairment we performed the cell viability MTT test 24h post-treatment and considered the increase with regard to the value obtained before treatment (Fig. 3). In this figure zero for the Y axis means therefore no change with a same number of cells that remained in a stable over 24h, positive value means proliferation and decrease cell death. This was made in the reference state (no addition or vehicle), in presence of CCCP at the level for 100% interference with OXPHOS and in presence of oligomycin as the reference state with no OXPHOS. The potency of rescue by glycolysis was made variable by changes in the concentration of glucose (1, 3, 25 mM) and set to minimal (none) if glucose was replaced by galactose. Increasing glucose concentration in absence of drugs had no significant effect (Vehicle: top left in Fig. 3). In contrast, when the ATP synthesis by mitochondria was cancelled (oligomycin) or impacted with CCCP at a level of 100% of decrease in OXPHOS, the cell fate was linked to glucose concentration. With 1mM glucose cell death occurred, increasing glucose to 3mM restored energy homeostasis and maintained survival, and 25 mM of glucose appeared able to promote some proliferation. In presence of galactose some growth was observed in the control conditions but massive cell death was observed when OXPHOS was suppressed by Oligomycin or with the 100% interference caused by CCCP. This result illustrates that galactose renders these cells fully dependent on OXPHOS.

**Figure 3:**
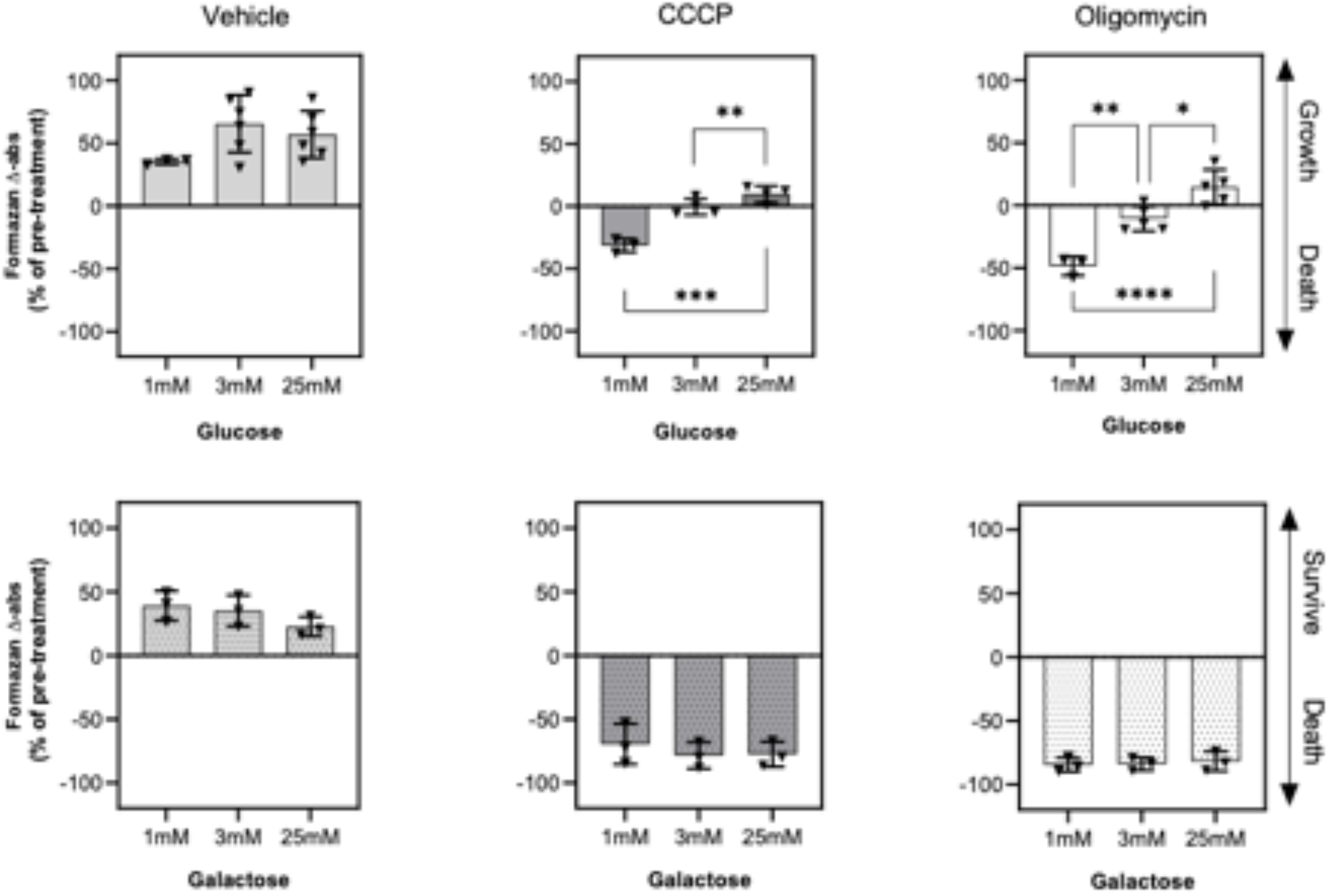
Effect of mitochondrial ATP abolishment on the cellular viability and growth. Cell viability and growth were determined by MTT assay at 24h post-treatment. Data shown are the difference in % of pre-treatment values. Data are expressed as well as the mean value ± SD of at least three independent experiments. * = p < 0.05; ** = p < 0.01; *** = p < 0.001; **** = p < 0.0001

The profiles observed with CCCP and oligomycin (reference state with zero OXPHOS) were very similar, then full inhibition of the mitochondrial OXPHOS enzyme by oligomycin or maximal interference with OXPHOs with the uncoupler CCCP had the same influence on cell viability/growth. Hence although the complete suppression of OXPHOS by uncoupler could not be demonstrated, the difference between oligomycin addition and 100% interference with uncoupler, if any, was of marginal importance with glucose concentrations of 3 or 25mM and gave confidence in our estimation of the OXPHOS rate (Fig. 2 D-F).

### Impact of exposure to uncouplers (thiazolides, CCCP) on virus release

Then we evaluated how a decrease in OXPHOS caused by uncouplers impacts on viral production. The phoenix ECO cells were transfected with a retroviral vector for EGFP and the virus-producing cell monolayers grown in 3mM glucose were treated for 24h with different concentrations of TZ, RM, or CCCP to decrease by 25%, 50%, 75%, or 100% OXPHOS. Controls included no treatment, solvent (DMSO) and oligomycin (full suppression of OXPHOS). After the treatment period, the viral release in the medium was estimated from the number of viral RNA copies in the supernatant as assessed by RTqPCR (Fig. 4). Oligomycin lowered significantly viral RNA release revealing the importance of OXPHOS in this process. A same effect was observed with CCCP, RM or TZ when OXPHOS decrease was settled to 75 or 100%. For a significant part, it could result from impaired cell growth during this 24h period of viral production (Fig. 4). At the opposite milder interference (25-50%) had no effect on the viral RNA release. The infectious potency of the viruses released with the lower levels of interference (25, 50%) was then quantified, for this purpose the viral particles were concentrated by co-precipitation with calcium phosphate. This was made with the same media than those presented in Fig. 4. In addition to concentration, the precipitation step eliminated the uncouplers and thus prevented their possible impact on the infection test. Growing NIH-3T3 cells were then incubated with Ca-Virus co-precipitate for 24h. After this infection period, the NIH-3T3 cells were maintained in a new culture media for 72h and efficiency of infection assessed by detection of GFP fluorescence (Fig. 5A). This test revealed roughly a 50% decrease in infectivity with 25% percent of interference with cellular OXPHOS. This effect did not appear amplified further with 50% interference on OXPHOS. There are two possible interpretations: either a significant part of the viral particles detected on the ground of RNA content were not infectious at all, or the probability for each viral particle to achieve a successful infection was decreased by half.

**Figure 4:**
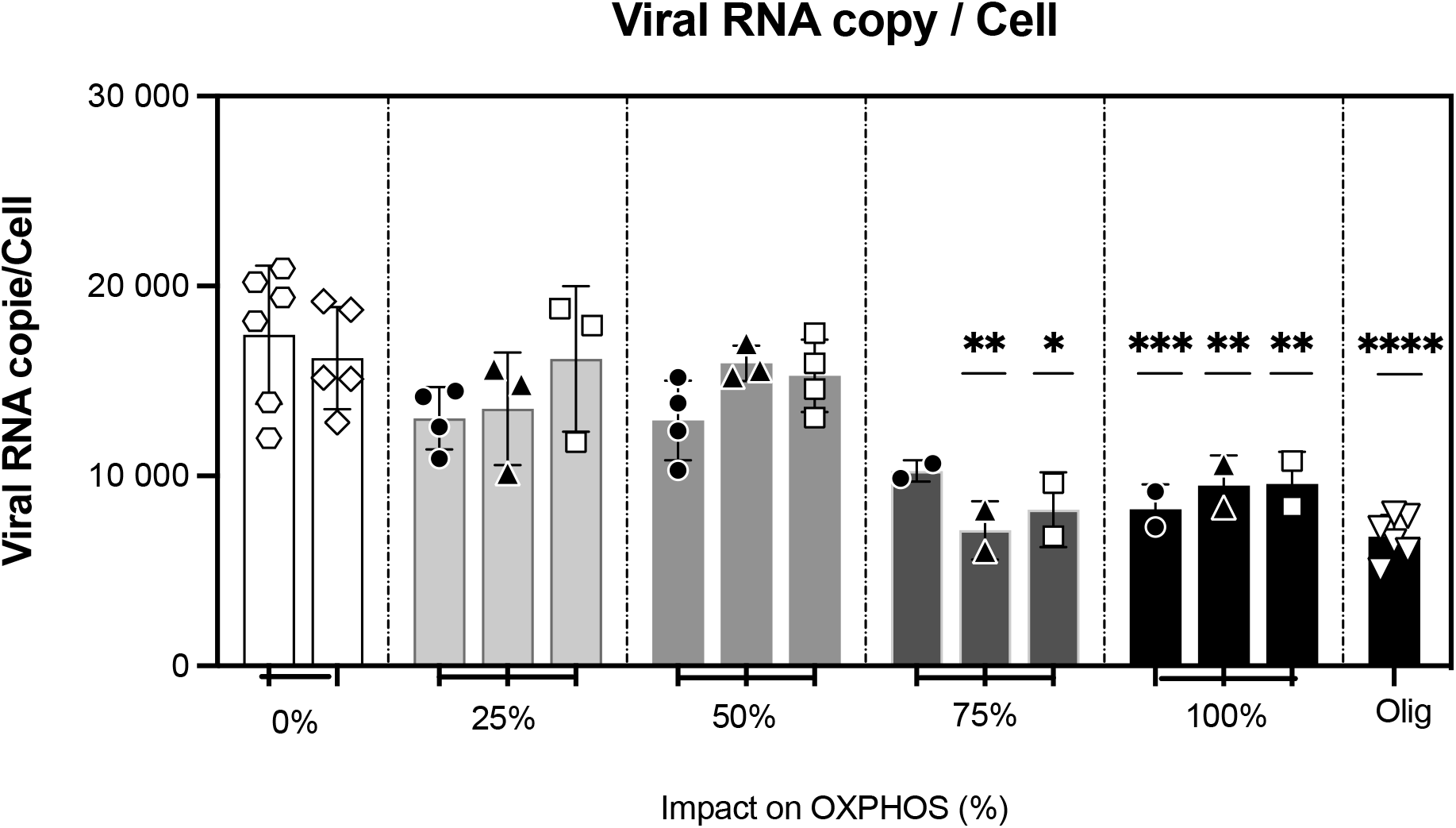
Increasing impact on OXPHOS and viral RNA release. The number of viral RNA copies in the supernatant was assessed by RTqPCR titration at 24h post-treatment. The levels of interference with OXPHOS are indicated below the groups of histograms and coded from black (0% interference) to white (100% interference and full inhibition by oligomicin). The reference data point: hexagons for control (no addition) and diamonds for vehicle (DMSO), mitochondrial poisons: squares, reference uncoupler (CCCP); downward triangles, oligomycin. Thiazolides are indicated by black symbols, circles, tizoxanide, triangles RM4848. Data shown are expressed as well as the mean value ± SD of at least two independent experiments. * = p < 0.05; ** = p < 0.01; *** = p < 0.001; **** = p < 0.0001, for comparison with vehicle.

**Figure 5:**
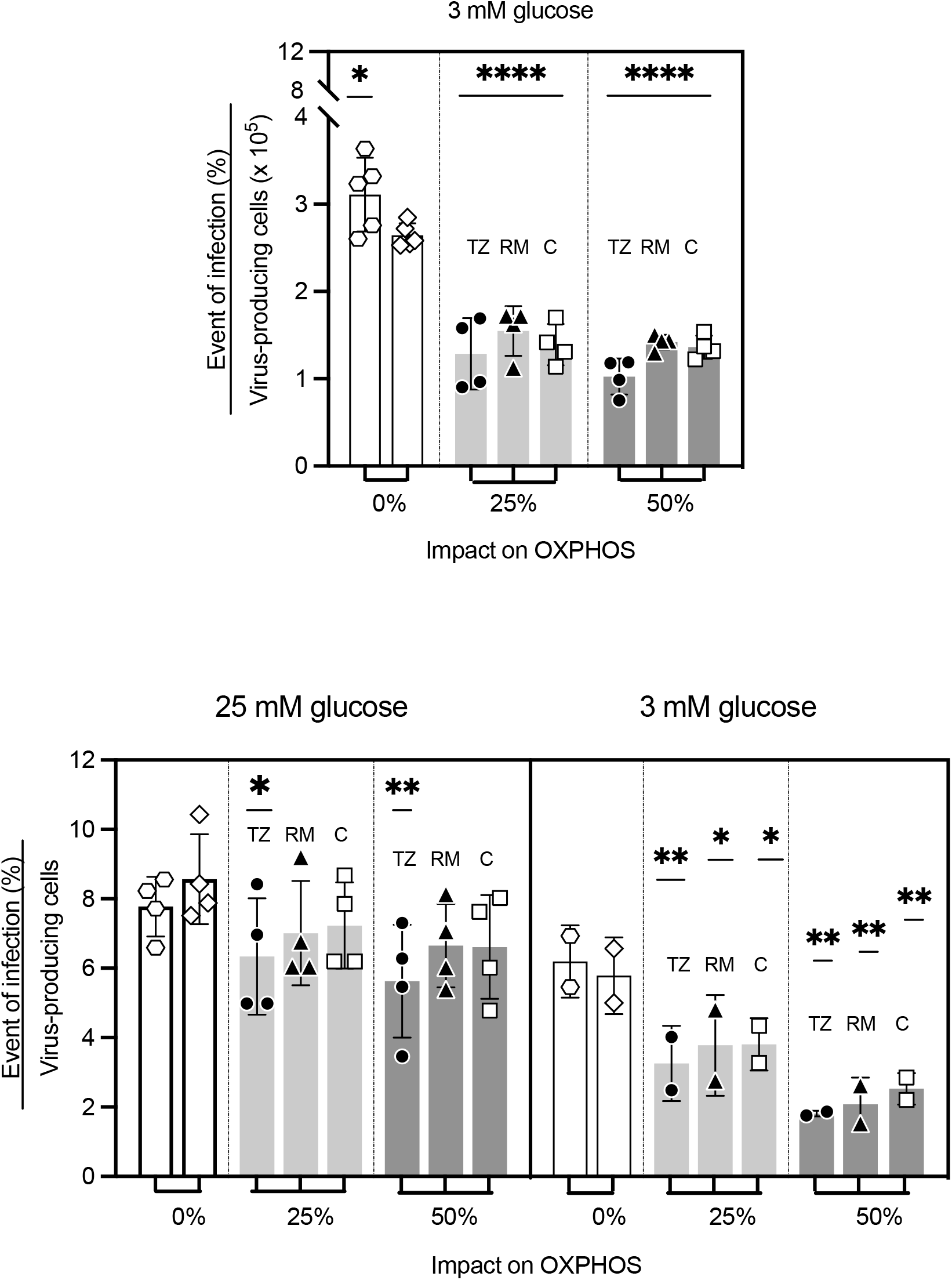
Increasing impact on OXPHOS and infectivity of newly produced viruses. Two series of experiments were performed. Top: experiments made with 3 mM Glucose in the medium and in which the viral RNA release was quantified (Fig. 4). Bottom left: experiments with 25 mM glucose in the medium, no quantification of viral RNA was attempted, in two of these experiments new data points with 3mM glucose were included (bottom right), rest of the legend as above.

To confirm that the lower quality of the collected viruses was indeed due to a bioenergetic disturbance, a new pool of Virus-producing cells cultured in a media with 3mM or 25mM glucose were treated for 24h with TZ, RM, or CCCP to decrease by 25% or 50% the OCR for ATP synthase (Fig. 5B). In these experiments we replicated the results presented in Fig. 5A and, in contrast with above, with 3mM glucose a dose response effect of the uncouplers appeared detectable. Increasing glucose in the media to 25mM restored the infectivity to levels not significantly different than the controls (RM, CCCP) but apparently still slightly lower with TZ.

### Cellular adenine nucleotide energy charge

We undertook a detailed analysis of content in adenine nucleotides to evaluate presence of signs of cellular energy depletion at the milder levels of impact sufficient to cause a significant loss in in the viral experiments (Fig. 5). The measured cellular content in adenine nucleotides varied greatly from one experiment to another (not shown), but in contrast the adenine energy charge (AEC) that is given by the ratio: (ATP+^1^/_2_ADP)/ (ATP+ADP+AMP) appeared reproducible and consistent (Fig. 6). Impairment of OXPHOS with oligomycin caused a serious decrease in AEC with 3mM glucose while it had no effect with 25mM glucose (Fig. 6 black arrows). Whatever the glucose concentration, impact of the 25% or 50% levels on OXPHOS had no influence on the AEC when compared to control (DMSO) with the exception of a possible marginal decrease (3mM glucose RM50%). Unchanged AEC does not imply that fluxes remain the same but means that consumption and production remained matched. Therefore, no signs of cellular energy depletion/imbalance were present at interference levels of OXPHOS able to impact seriously on viral particles infectious properties: 3mM glucose and 25 or 50% impact on OXPHOS.

**Figure 6:**
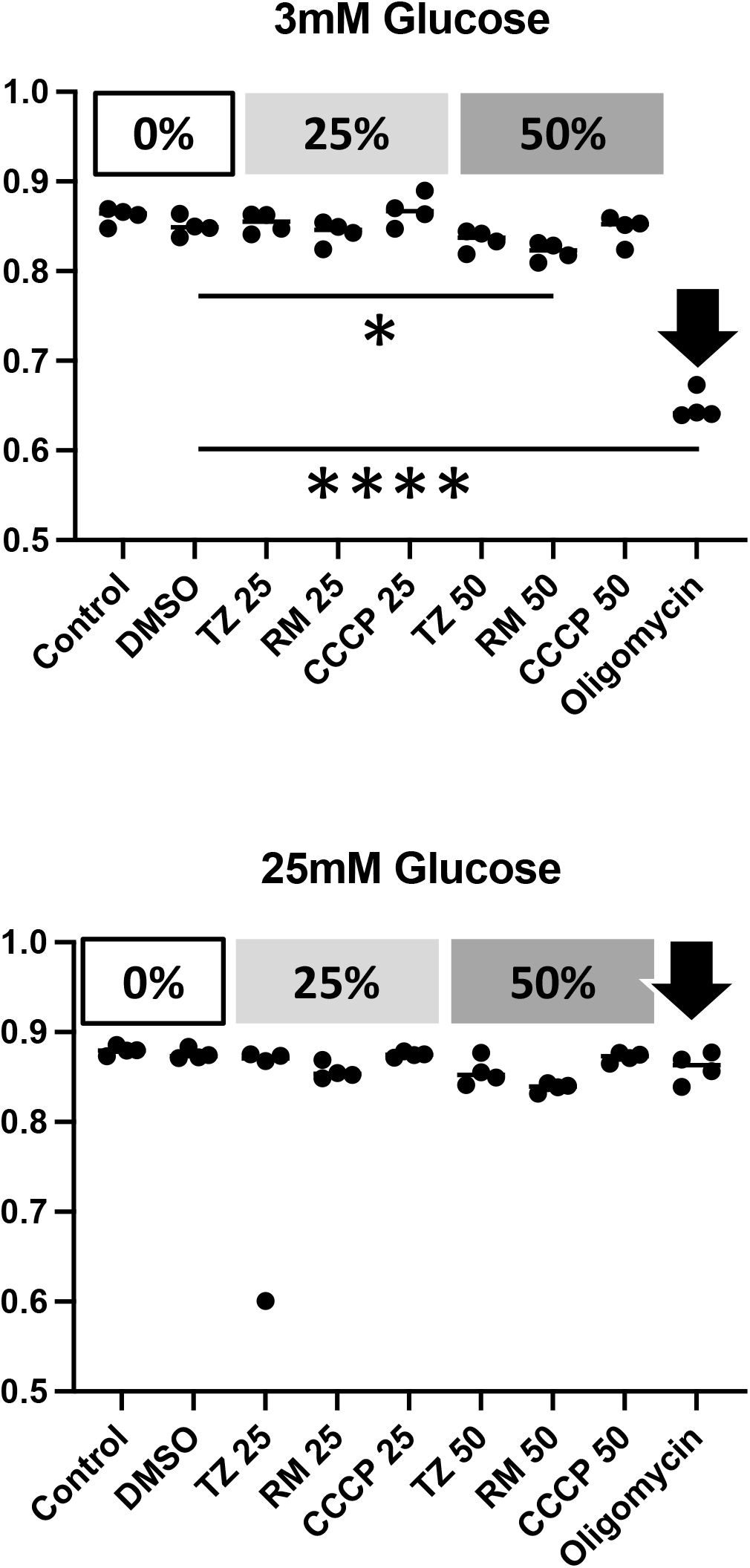
Adenine nucleotide energy charge (AEC). AEC is a dimensionless ratio given by the formula: (ATP+^1^/_2_ADP)/ (ATP+ADP+AMP), the Y ordinate starts at the value 0.5, which is the value of AEC obtained if ADP represents 100% of adenine nucleotides. The level of interference of drugs with OXPHOS are indicated above with the same grey code as in figure 5. The black arrow highlights the oligomycin reference with OXPHOS set to zero. Drugs with levels of interference with OXPHOS are indicated below X axis. Data point are shown with their median (four determinations/independent experiments).

### vMitochondrial studies

The uncoupling activity of TZ and RM was further characterized in isolated mitochondria. Freshly isolated rat liver mitochondria were incubated in mitochondrial respiration buffer, supplemented with rotenone (inhibitor of complex I) and oligomycin (inhibitor of ATP synthase). Respiration was stimulated with succinate (substrate of complex II) and oxygen consumption rate (OCR) and membrane potential (ΔΨ) were measured during successive additions of TZ, RM or CCCP. TZ, RM and CCCP caused an oligomycin insensitive dose-dependent increase of oxygen consumption (Fig. 7 A-C: solid line), coupled to a parallel decrease in mitochondrial membrane potential (ΔΨ) (Fig 7 A-C: dotted line). This observation tallies with increase of the passive permeability (conductance) of the inner membrane to protons pumped by the respiratory chain, the molecular event causative for uncoupling. Not too much attention should be paid to the apparent differences when membrane potential drops below 120mV (see M&M section). Supplementary experiments included direct comparison of effect of TZ or RM in presence/absence of CAT or Cyclosporin A. CAT is an inhibitor of the adenine nucleotide translocase (ANT) that exchanges ATP against ADP across the mitochondrial inner membrane. CSA is an inhibitor of the mitochondrial transition pore (mPTP). None of them modified the uncoupling activity suggesting no interaction of thiazolides with the ANT or mPTP. Therefore, although the number of possible other interactions is immense, these results eliminated two important actors of the mitochondrial inner membrane and possible causes for highly deleterious impact on cellular bioenergetics/survival. A direct comparison of the relative efficiency of thiazolides and CCCP is represented (Fig. 7D), which takes into account both OCR and membrane potential. The three compounds led to a similar maximal value for the product OCR×ΔΨ. The proton pumping by the respiratory chain is supposed to be directly proportional to OCR. Therefore, the product OCR×ΔΨ is linearly correlated to the power dissipated by the proton current in a circuit where the respiratory chain is a generator associated to a resistance (conductance of the inner membrane to protons) that is influenced by uncoupler dosage (inset in Fig. 7D).

**Fig. 7.**
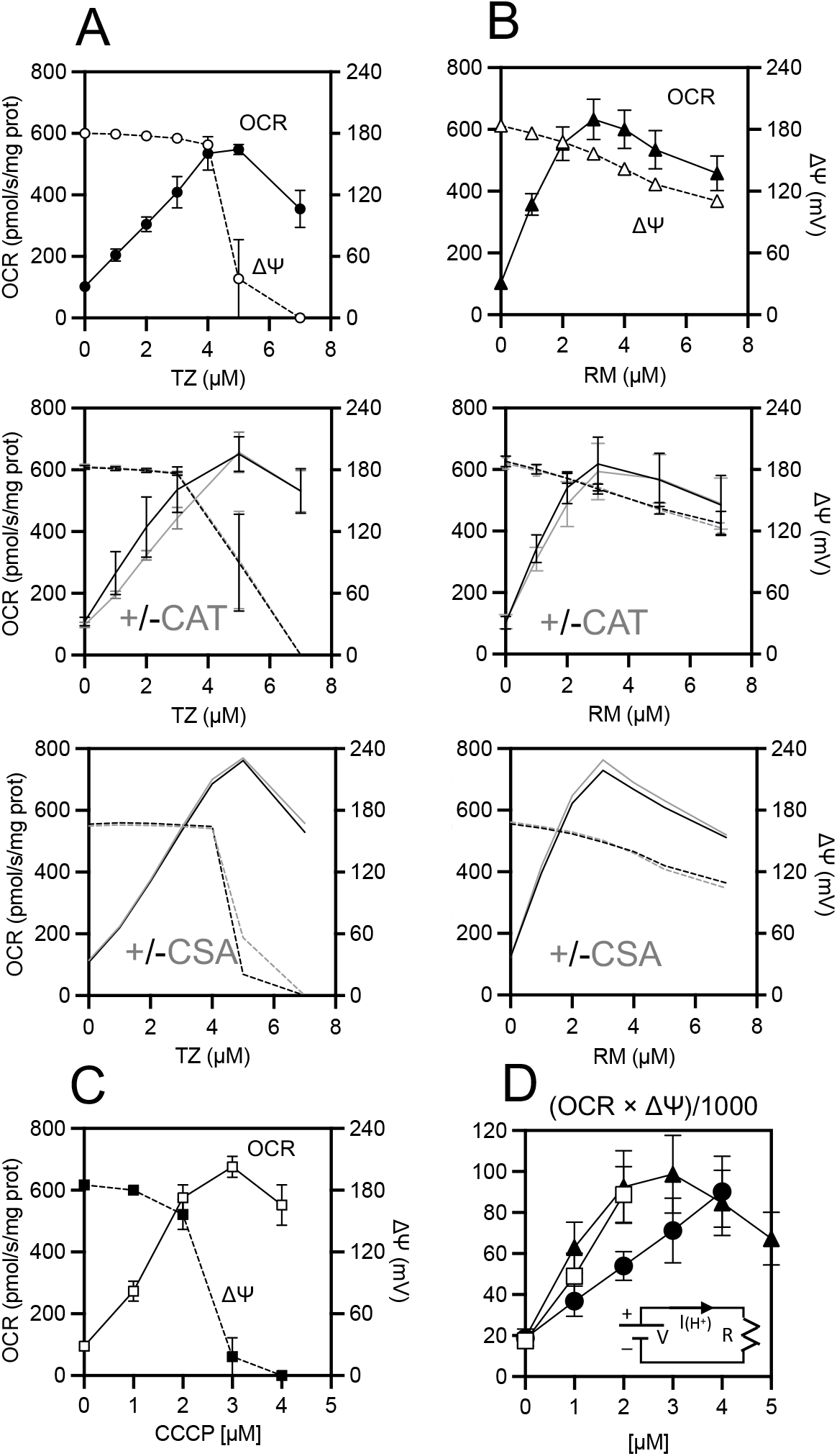
Mitochondrial uncoupling by TZ, RM and CCCP. ATP production (Oxphos). A-C: Dose response to thiazolides and to CCCP. The value of OCR (solid line) and of mitochondrial membrane potential (milliVolts) are shown starting from the oligomycin, non-phosphorylating rate (X=0) and with increasing concentration of TZ (A), RM (B) or CCCP (C) with a different range of concentrations. Experiments comparing thiazolides effect in presence (grey lines) or absence (black lines) of CAT or CSA are shown without symbols for clarity (CAT N=3) CSA (single experiment). D: product of OCR and membrane potential for the three uncouplers symbols as in A-C, for TZ and CCCP values are shown up to the maximal value and not for higher concentrations.

We then examined the consequences of thiazolides on mitochondrial OXPHOS with mitochondria respiring in presence of succinate and in absence of oligomycin. Addition of a saturating dose (1-2mM) of ADP caused immediate decrease in membrane potential accompanied by increase in OCR with establishment of a new steady state for mitochondrial respiration with a maximal OCR rate. The relationships between membrane potential (X axis in Fig 8A) and OCR (Y axis in Fig 8A) in the different experiments are represented for these two steady states (grey diamonds) designated with P (phosphorylating) and L (Leak) a term to highlight that OCR in this state is explained by endogenous proton leakage through the mitochondrial inner membrane. Increasing concentrations of the uncouplers under the non-phosphorylating conditions (see above) caused a progressive transition from L to P ordinates. Therefore, with regard to OCR and ΔΨ the maximal uncoupled state caused by TZ, RM or CCCP cannot be distinguished from the maximal rate of oxidative phosphorylation. While this illustrate that uncouplers would substitute to ADP with regard to effect on mitochondrial respiration, the interference between both remained to be examined. Therefore, we used the luminometric assay for ATP after kinetic experiments (Fig. 8B) to compare the real ATP production rate in absence/presence of TZ or RM (Fig 8B). Importantly, succinate oxidation in presence of rotenone prevented the non-OXPHOS ATP production by succinyl-Co-A ligase (EC 6.2.1.4). The amount of ATP produced could then be compared to the oxygen consumed over a same period of time. Their ratio (ATP/O) quantifies the efficiency of conversion of the redox energy released during succinate oxidation into ATP. The value for this ratio was 1.8 in absence of drugs almost equal to the expected theoretical value indicating good mitochondrial preparations and was lowered to 1 in presence of concentrations of RM or TZ that lowered the mitochondrial membrane potential to the same extent as observed with ADP addition (TZ RM in fig. 8C). With a concentration decreased to the half (Half TZ/RM in Fig. 8C), an intermediate value was obtained. As expected, the effect of TZ or RM on mitochondrial OXPHOS was dose dependent. Studies with cells had to rely on another estimation of the OXPHOS rate supposed to be directly proportional to the difference between OCRs in absence/presence of oligomycin (OXPHOS OCR). In these mitochondrial studies these two modes of calculation could be compared in presence/absence of TZ or RM (Fig. 8D). This revealed a linear trend but interception for X=zero (no more oligomycin sensitive OCR) did not coincide with the zero ordinate for ATP formation rate (Y axis). Then, full uncoupling as judged from values of ΔΨ (Fig. 8C) or OCR (Fig. 8D) did not annihilate OXPHOS and consequently the difference between OCRs observed in presence/absence of oligomycin appeared linearly related to the OXPHOS rate but would underestimate it.

**Fig. 8.**
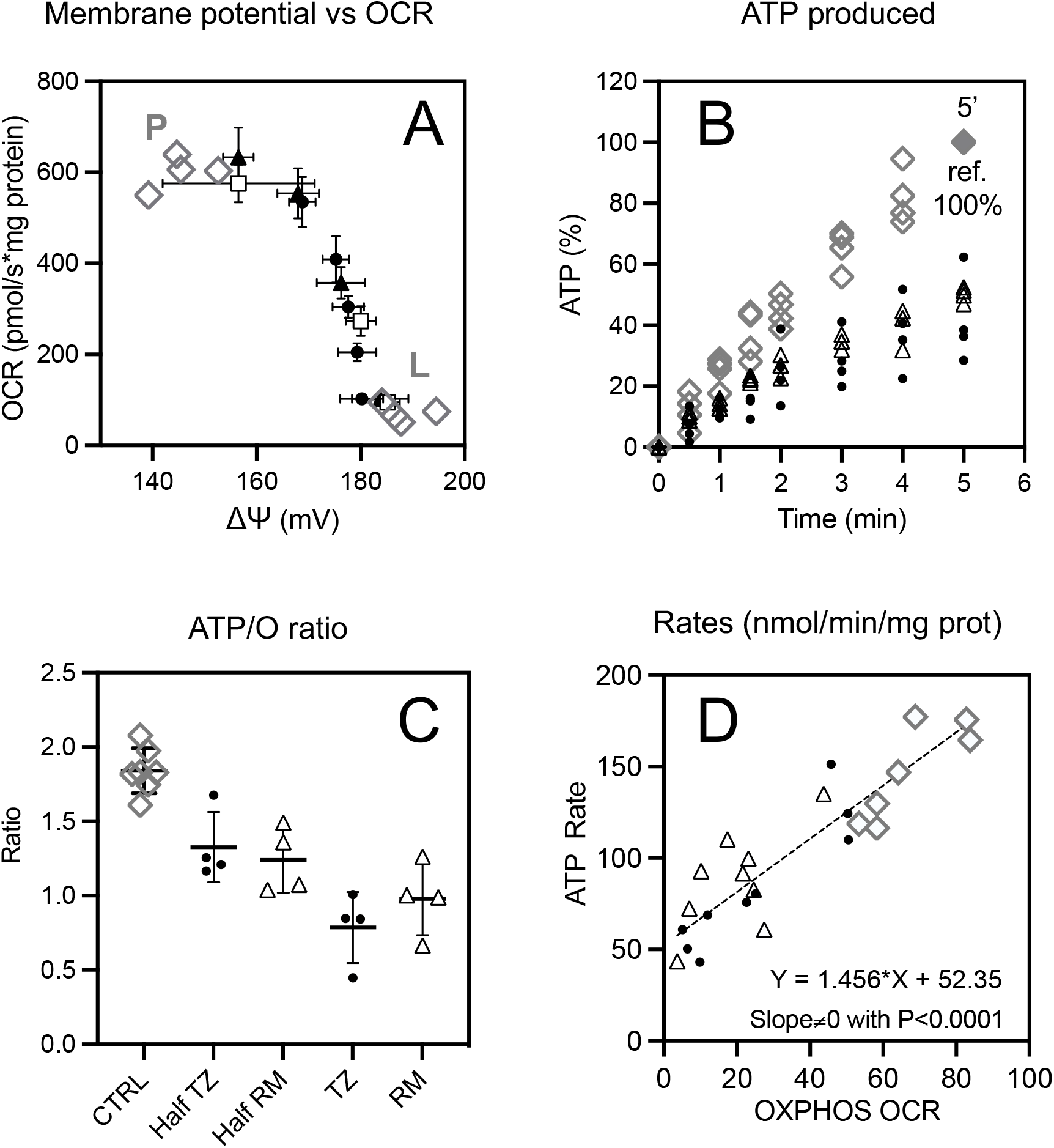
Mitochondrial ATP production (Oxphos). (A) relationship between oxygen consumption rate and membrane potential in mitochondria at steady state with individual data points (grey diamonds) shown for different experiments (independent mitochondrial preparations), “P” indicates the phosphorylating state (ADP present) while L indicates the values in absence of phosphorylation (oligomycin present). The effect of different concentrations of thiazolides (filled black symbols) with circles (TZ) and triangles (RM) and of CCCP (empty squares) are shown. (B) ATP sampling during mitochondrial respiration and dosage by bioluminescence, expressed in relative units (100% is the ATP produced by the respective control in five minutes), symbols: control (grey diamonds), TZ (black circles) or RM (empty triangles), at concentrations resulting in the same depolarization than ADP addition. (C) ATP/O ratio for control (CTRL) and in presence of TZ or RM at the concentration used in B or decreased by half, individual values are shown with their mean and error bars (sem). (D): Relationship between ATP synthesis rate obtained from ATP dosage and from the decrease in OCR caused by the ATP synthase inhibitor oligomycin. Individual values shown in B, C, D were obtained in at least three independent experiments (different mitochondrial preparations).

## Discussion

### Viruses are dependent on cellular bioenergetics

A shared property of viruses is their dependence on their host cell metabolism, assembly of the virus particle implies a large number of different energy demanding steps. For example, the polymerization of viral nucleic acid corresponds to one ATP per base, incorporation of one amino acid into a viral protein to four. In addition, maturation steps with post translational modifications and viral assembly processes are other consumers of cellular energy (ATP). Virus assembly constitutes therefore a supplementary burden for cellular energy metabolism and a competitor with normal cellular processes for the access to cellular ATP. If production of viral particles would compromise the balance between ATP generation and consumption this would cause cell death and end viral replication process. Accordingly, virus propagation depends also on conformity with the constraints of cellular bioenergetics. Consistent with this, several viruses have been shown to stimulate cellular metabolism,^22 23 24^ and this is thought to prepare the cell to the increase in ATP demand caused by viral replication. Here we explore how the viral replication is altered when cellular bioenergetics is challenged by adverse conditions. Our starting hypothesis was that the wide antiviral properties of nitazoxanide (NTZ) result from its uncoupling activity and therefore could be exerted on any scheme for viral production hence in the convenient model used here, remote from those used so far to evidence NTZ antiviral properties.

### Interference with mitochondrial bioenergetics explains antiviral effect of NTZ

A prediction directly derived from this hypothesis was that the effect of nitazoxanide (NTZ) and that of an uncoupler of mitochondrial respiration on viral replication would be identical and the dose response correlated to the intensity of the uncoupling effect of these two different molecules. To this purpose we compared the antiviral effects of TZ the active form of NTZ with a reference mitochondrial uncoupler (CCCP), and with another thiazolide differing by replacement of the nitro group (RM). The first step was to calibrate their respective effect on the cellular model of interest to generate identical levels of uncoupling by the three molecules (Fig. 2). Then the impact of these levels of OXPHOS uncoupling on viral release were examined and the results were entirely consistent with expectations because the antiviral effects of TZ, RM and of the uncoupler CCCP were the same when concentrations of these different molecules were adjusted for a same impact on mitochondrial bioenergetics *in cellulo*. This was true both for RNA release (Fig. 4) and infectivity (Fig. 5). Accordingly, the simplest explanation would be the following: what matters is the intensity of the uncoupling effect and not the nature of the agent and then the antiviral activity of TZ, in this model, is consequence of its uncoupling activity. A mode of action of NTZ based on the inhibition of pyruvate metabolism was proposed,^3^ in mammalian cells the target enzyme would be the pyruvate dehydrogenase (PDH). Since the three uncouplers inhibited viral replication to the same extent effect on PDH should be either not required or supposed to be exerted with a same intensity by the three uncouplers. This effect on PDH is thought to be dependent on the NO_2_ group of TZ it would then be absent with RM (Fig. 1) and moreover a similar effect on PDH is hardly conceivable for CCCP. Then, if present, effect on PDH appears of negligible importance here and indications for it would require different experimental schemes.

### Mild uncoupling shows significant antiviral effect

To be an acceptable explanation for the antiviral effect of NTZ *in vivo* the level of uncoupling required for repression of virus propagation should be more deleterious for the virus than for its host. Full uncoupling (100% of interference with OXPHOS) is feasible *in vitro* because substrate linked phosphorylation steps are not sensitive to uncouplers and consequently intense glycolysis could substitute to OXPHOS (Fig. 6). However, this makes no sense *in vivo* and to be an acceptable explanation low levels of interference should exert adverse effect on viral propagation. The second tenet was therefore that the viral replication cycle is exquisitely sensitive to bioenergetic challenge because cells undergoing energy shortage re-orient and reduce the ATP use according to priorities^20 21^ and the production of new viral particles would not be of high priority. The usual challenge for animal bioenergetics (OXPHOS) are lack of oxygen or of substrates hence associated to reduction in the flux of energy metabolism and hypothermia.^20 21^ In contrast, the cellular response to uncoupling is to increase substrate and oxygen use to compensate for the lower yield of OXPHOS and this was expected to result also in re-orientation of ATP use to preserve balanced ATP production and consumption rates. Our results are fully compatible with these hypotheses. Firstly, under the lowest level of interference used here (25% uncoupling) the preservation of AEC indicated that virus producing cells could keep balanced ATP generation and consumption rates. Secondly, these adaptations impacted significantly on virus infectivity that was decreased by half suggesting that even lower levels of interference would be effective.

### Mild uncoupling impacts selectively virus maturation

However, not all steps of viral replication were concerned by this repression under mild bioenergetic challenge: the repression of RNA release required strong interference on cellular bioenergetics (75% 100% uncoupling). In this respect the use of two different criteria for virus release was of importance as the lack of infectivity of viral particles released as “packaged RNA” and produced in same amount as in control excluded a mere toxic effect on the virus producing cells. Accordingly, a process of viral formation subsequent to viral RNA packaging is expected to be impacted by mild bioenergetic interference. It tallies with other studies made with influenza^25^ or SARS-Cov2 viruses^19^ in which NTZ impaired maturation of viral particles. The strategy used here for comparison of TZ with an uncoupler could well be transposed to experimental schemes adapted to pathogenic viruses. An unresolved issue is whether defect in maturation impacts equally all viral particles or if at the opposite a part is excluded from the maturation process. Few more issues may deserve consideration: 1) does bioenergetic interference ground antiviral properties of other drugs? 2) Does this pharmacological intervention reproduce/reveal characteristics of virus host interaction? 3) Why the uncoupler NTZ could be a safe molecule while the previous use of the uncoupler dinitrophenol showed severe adverse effects.

### Cellular response to bioenergetic challenge could explain unexpected antiviral effect

The impact on virus maturation/infectivity is expected to result from a general response with aim to prevent a deadly imbalance between energy (ATP) supply and use.^20 21^ It could therefore be triggered by drugs impacting directly on energy metabolism and by others that by any means increase cellular energy demand. This may explain unexpected antiviral properties observed with a significant number of molecules. This is often observed *in vitro* and the issue is whether or not this applies *in vivo*. In this respect, concentrations of TZ found in patients treated with this drug (7-100µM) range well within the values used here. However, at least two supplemental factors are to be considered: Firstly, binding of TZ to plasma proteins *in vivo* would decrease its availability but at the opposite lipophilicity is expected to drive TZ towards membranes (the target site for uncoupling effect in mitochondria). Secondly, our experiments were made under conditions where oxygen supply was not supposed to be limiting, this might be different *in vivo*. Uncoupling deteriorates the yield in ATP per oxygen (ATP/O ratio) and under conditions of oxygen limitation even a modest decrease in ATP/O ratio has serious consequences on the balance between ATP consumption and production.^26^

### Host-virus interactions

A recurring motif in biomedical research is that therapeutic strategies reflect and sometimes reveal physiological mechanisms mobilized for related purposes. Fever results from an increased heat production and therefore increases energy expenditure as uncoupling does. Re-routing of ATP according to priorities was documented with hypometabolic states, which ensure adaptations to oxygen deprivation and improve also resistance to starvation. Then while fever (hypermetabolism) and anorexia appear contradictory responses to infection, both would converge to trigger cellular bioenergetic responses adverse to viral propagation. It would be difficult for a virus to become resistant to energy shortage and overriding cellular priorities would increase the risk of premature cell death and interruption of virus replicative cycle. A virus adaptive strategy would be to promote cellular energy metabolism, this has been observed.^22 23 24^ In the present study the viral particles (packaged RNA) were produced in normal quantities but globally their infectivity was reduced. This would preserve the host/virus duo: On one side the host would be protected from a deadly viral propagation and on the other side the host immunity would still be diverted by still numerous “fake viruses” increasing the chances for the virus to escape to complete destruction.

### Nitazoxanide a safe uncoupler used for years

A final point to be examined is whether or not the level of uncoupling as observed here is compatible with NTZ be a safe drug and this because former use of the uncoupler dinitrophenol proved to generate severe side effects. Firstly, the aim was different past use of dinitrophenol had the purpose of increasing energy expenditure for weight loss, the higher the increase the faster/greater was the weight loss, with risk of too high dosage. Secondly, it could rely on a different mechanism for toxicity than uncoupling. Two lines of evidence would support this contention: 1) Dinitrophenol is a poorly efficient uncoupler and dosage needs therefore intense exposure to this phenol derived compound leaving place for other toxic mechanisms to develop. 2) A potentially harmful side effect of dinitrophenol would result from the fact that several uncouplers, and notably dinitrophenol, interact with the ATP/ADP exchanger of the mitochondrial inner membrane.^27^ Then in presence of dinitrophenol the mitochondrial ATP would be produced with a lower yield because of uncoupling and furthermore ATP export from mitochondria to the rest of the cell would be impaired. This dual hit may well increase greatly toxicity. Our studies on the effect of TZ and RM effect on isolated mitochondria excluded interaction with the ATP/ADP exchanger. Similarly, we excluded another possible route of toxicity: the interaction with the mitochondrial transition pore, a trigger of apoptotic cell death. Finally, we should consider the existing knowledge about uncoupling accumulated after (independently from) the “dinitrophenol accident” for recent reviews see.^28 29^ Uncoupling increases energy expenditure and at the level of the organism might be considered^28^ or not^30^ to improve metabolic status. In addition the concept of “mild-uncoupling” reflecting a modest impact on the coupling state of mitochondria and on energy expenditure is considered as a mean to mitigate the mitochondrial generation of reactive oxygen species hence to prevent oxidative stress but this remains controversial.^31 32 33^ Hence while re-routing of energy is our hypothesis, impact on viral maturation through modulation of ROS generation and or cellular redox state might be an alternate explanation. The rescue by high glucose of the effect of TZ does not support this alternate hypothesis and led us to the conclusion that energy supply (ATP) was the relevant issue here. One or the other would not change that the mode of action of NTZ would rely on its effect on mitochondrial bioenergetics and be an example of use of a safe uncoupler, or of an uncoupler in the safe range. For this reason, further therapeutical properties of NTZ should be scrutinized.

## Materials and Methods

### Drugs and reagents

Tizoxanide (dAcetyl nitazoxanide) and RM4848 were provided by Romark Laboratories, 2-[2-(3-chlorophenyl)hydrazinylyidene]propanedinitrile (CCCP) (Sigma-Aldrich), rhodamine 123, cyclosporine A, or carboxyatractylate were dissolved in dimethyl sulfoxide (DMSO); rotenone and oligomycin were dissolved in 1:1 (v/v) DMSO:Ethanol; cyanide was dissolved in water.

### Cell culture and treatments

Phoenix Ecotropic (ECO) (ATCC – CRL-3214™), and NIH-3T3 (ATCC – CRL-1658™) were grown at 37 °C, with 5% CO2 in DMEM, high glucose, pyruvate supplemented with 10% FCS (Gibco – 31966021). 80% confluent cells were harvested by addition of trypsin-EDTA (0.25%), centrifuged, and resuspended in a new culture media (DMEM with desired concentration of glucose, containing 10% FCS). The same growth conditions were used for maintenance of transfected Phoenix-ECO cells (the virus-producing cells). Growth of Phoenix ECO cells, viral particle production and infection of NIH-3T3 required L2 confinement levels and experiments were made under the authorization of local authorities (N°1878). In experiments using custom concentrations of glucose, a mix of DMEM high glucose (Gibco –31966021) and DMEM no glucose (Gibco – 11966025) has been performed. DMEM 1mM galactose was obtained by mixing a corresponding volume from 1M galactose stock solution and DMEM no glucose. Phoenix-ECO/virus-producing cells were grown to reach around 40-50% of confluence at 24 of culture incubation. Treatments of the cell monolayers were performed by adding a volume of the relevant media that contain drugs at desired concentration and kept in the culture medium for the entire time of the experiment (24h), unless differently specified. Controls received equal amounts of vehicle (DMSO).

### Cell transfection and selection of a virus-producing cell population

Phoenix-ECO (ATCC® CRL-3214™) were transfected with pMMLV[Exp]-EGFP/Puro (VectorBuilder, VB010000-9307ddn) using jetPRIME®transfection reagent (Polyplus). After few days of puromycin selection, Phoenix GFP were sorted by flow cytometry to obtain 3 cell populations: Phoenix GFP-, GFP+, GFP++. This last population had the highest level of fluorescence and was used as the virus-producing cells model.

### Cell survival and proliferation assay

The MTT assay is a convenient method to quantitate both Cell survival and proliferation because it quantitates the number of living cells. It is based on the capacity of viable/active cells to reduce the 3-(4,5-dimethylthiazol-2-yl)-2,5-diphenyltetrazolium bromide (MTT) to formazan. Cell survival and growth were appreciated by the determination of the increase in formazan formation obtained in 24h post-treatment. Zero means therefore no change with viable cells that remain stable, positive value means proliferation and decrease cell death. This was made in the reference state (no addition or vehicle).

According to the cell culture protocol, equal number of cells (15.000/well for culture in glucose or 22.000/well for culture in galactose) were seeded in two different 48 well plates and cultured in DMEM without phenol red, supplemented with custom concentrations of glucose or galactose. After 24h, one plate was used for MTT assay. It is the pre-treatment condition. For the second plate, d’AcNTZA, RM4848, CCCP or oligomycin were added in triplicate at desired concentration and kept in the culture medium for the entire time of the experiment (24h). Controls received equal amounts of vehicle (DMSO). After 24h of treatment, MTT assay for treated-untreated cells was performed.

MTT assay was performed as following: 1:3 v/v per well of new medium containing MTT substrate 50 ng/µl final was added to treated-untreated cells, and incubated for 4 hours at 37 °C, with 5% CO_2_ atmosphere. After this period, the formed formazan was extracted from cells and dissolved by adding 1:1 (v/v) of 2X (SDS 10% and HCl 0.01%) solution. After 24h incubation in room temperature, the quantity of formazan was measured by recording changes in absorbance (Δ-abs) at 570 nm (wavelength for maximum absorbance of the formazan) and 680 nm (reference wavelength) in an ELISA microplate reader.

### Viral RNA extraction and quantification

1.8.10^5^ of virus-producing cells per well were seeded in a 12-plate and cultured/ according to the culture protocol. After 24h, the cell monolayers were treated by changing the old media by one ml of a new media containing desired concentrations of dAcNTZA, RM4848, CCCP or oligomycin. Controls received equal amounts of vehicle (DMSO). At 24h post-treatment, the RNAs of 150µl of the supernatant were extracted and quantified using Retro-X™ qRT-PCR Titration Kit (Takara, cat# 631453), according to the manufacturer instruction.

### Precipitation of viral particles from the supernatant

3.6.10^5^ of virus-producing cells per well were seeded in a 6-plate and cultured according to the culture protocol. After 24h, the cell monolayers were treated by changing the old media by a 1.5 ml of a new media containing desired concentrations of dAcNTZA, RM4848, CCCP or oligomycin. Controls received equal amounts of vehicle (DMSO). At 24h post-treatment, after centrifugation of the supernatant 5 min at 1000g to eliminate cell debris, 1.2 ml from the supernatant were transferred in a new tube containing CaCl_2_ (8mM final) and incubated at room temperature for 45 min. After this period, Ca-Virus particles was precipitated from the supernatant by centrifugation 45 min at 16000g. A final step of the procedure consisting to elimination the maximum of the supernatant and resuspension of Ca-Virus co-precipitates in 400µl of DMEM high glucose, supplemented with 10% FCS and polybrene (8µg/ml final) was performed.

### NIH/3T3 cell infection

24h before viral precipitation, 3600 of NIH/3T3 cells per well were seeded in a 48-plate and cultured according to the culture protocol. Immediately after viral precipitation process, old media was removed from wells and NIH-3T3 cells were incubated with 200µl/well of Ca-Virus co-precipitate for 24h. After 24h post-infection the viral inoculum was removed and NIH-3T3 cells were maintained in a new culture media. Infection yield was determined at 72h post-infection by counting GFP fluorescence positive cells detection with a flow cytometer (Accuri C6).

### Extraction of cellular metabolites and measurement of adenine nucleotides

Cellular extracts were prepared by an ethanol extraction method^34^. Metabolite extraction had been performed in the conditions used for precipitation of viral particles from the supernatant. Briefly, 1 ml per well of Ethanol/Hepes 10m M pH 7.2 (4/1) was added to the treated/untreated cell monolayers and incubated for 3 min at room temperature. Cellular extract was then transferred in a new tube and incubated at 80°C for 3 min. The mixture was cooled down on ice and the ethanol/Hepes solution was eliminated by evaporation using a rotavapor apparatus. The residue was suspended in sterile water at 2.10^3^ cell/µl. Insoluble particles were eliminated by centrifugation for 10 min at 21 000 g and 4°C and the supernatant was centrifuged in the same conditions for 60 min. Metabolites separation was performed on an ICS3000 chromatography station (Dionex, Sunnyvale, USA) using a Carbopac PA1 column (250 × 2 mm; Thermo Electron) with the 50 to 800 mM acetate gradient in NaOH 50 mM described in^35^. ATP, ADP and AMP content were inferred from standard curves using pure compounds. AXP content corresponds to the sum of ATP + ADP + AMP contents. Adenylate Energy Charge was defined as AEC = (ATP + ½ ADP) / AXP)^36^.

### Mitochondrial preparation

Rat liver mitochondria were obtained from male 5-week-old-SPF Wistar rats (Janvier Labs). The liver was homogenized in a mitochondrial preparation buffer (300 mM sucrose, 5 mM tris base, 1 mM EGTA, pH 7.4) just after the animal was sacrificed (Authorization number B75-14-02). Mitochondria were isolated by 2 differentials centrifugations. The final mitochondrial pellet was suspended in the same buffer (about 50 mg protein/ml final concentration). Protein concentration was quantified by the BCA method with bovine serum albumin as a standard. The Respiratory control ratio (RCR) was used to assess the integrity of mitochondrial compartments. It is calculated by the ratio of oxygen consumption respiration at phosphorylating state and non-phosphorylating state (see OCR measurement). Usually, the RCR for a fresh preparation should be higher than 10. For all experiments, an RCR>13 was fixed as the threshold for acceptability of mitochondrial preparation.

### Measurement of oxygen consumption

The O2k-FluoRespirometer was used to evaluate the oxygen consumption rate (OCR) of isolated mitochondria and whole cells. This apparatus contains two chambers for OCR measurement, allowing a paired experiment between Treated and reference. Before experiment, the oxygraph O2k was calibrated and maintained at the desired temperature (25°C for isolated mitochondria, or 37°C for whole cells).

### Experiments with cells

Phoenix Ecotropic (ECO) cells 1,5.10^6^ cell/ml were transferred in each of the two chambers of the oxygraph O2k. After 5min period for equilibration, both chambers were closed to start OCR measurement, the OCR at this stage includes mitochondrial contribution with phosphorylation of ATP (Phos-OCR). After 5min, oligomycin (2µM final) was added in one chamber this causes complete inhibition of the mitochondrial complex V (ATP synthase) and therefore mitochondria are settled to non-phosphorylating conditions (Non-Phos-OCR). Then gradual additions of dAcNTZA, RM4848, or CCCP were performed until the maximal stimulation of respiration. At the end of the experiment cyanide (1μM final) was added in each chamber. Cyanide is an inhibitor of mitochondrial respiration and the OCR observed in presence of cyanide quantitates the “non-mitochondrial OCR”. In our experiments it was insignificant, and for this reason was not considered further.

The decrease in OCR caused by oligomycin will be considered hereafter as a quantitative evaluation of the rate of ATP regeneration by OXPHOS, and will name this difference of the OCR before and after addition of oligomycin “OCR for ATP Synthase”. This OCR for ATP Synthase as observed following additions of the drugs (dAcNTZA, RM4848, or CCCP) was then expressed in percent of its reference value in absence of drugs (basal) as follow:

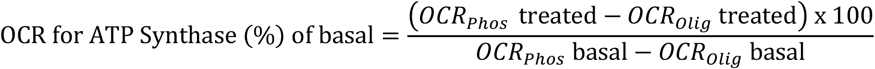

### Experiments with isolated mitochondria

Mitochondria were resuspended in the mitochondrial respiration buffer (KCl 100mM, Sucrose 40mM, TES 10mM, MgCl_2_ 5mM, EGTA 1mM, Phosphate K 10mM, fatty acid free BSA 0.2%, pH 7.2) at a concentration of 0.5 mg protein/ml final and distributed in each chamber of the oxygraph O2k. After 1min period for equilibration, both chambers were closed. For RCR determination, respiration at phosphorylating state was obtained by following additions of glutamate/malate (5 mM) and ADP (1.25 mM). After stabilisation of the OCR, oligomycin (1μM) a specific inhibitor of ATP synthase was added to measure respiration at non-phosphorylating state.

For experiments, 7.5mM of succinate, 5μM of rotenone (inhibitor of complex I) and 1μM of oligomycin were added to mitochondria to induce respiration at non-phosphorylating state (basal). According the experiment aims, gradual additions of different drugs were performed. At the end of the experiment cyanide (1μM final) was added in each chamber to quantify the “non-mitochondrial OCR”.

The membrane potential (ΔΨ) was evaluated by using the probe Rhodamine 123, for all the ΔΨ evaluations shown here the Rhodamine 123 fluorescence was measured with fluorometric module of the O2K and was synchronous to OCR recording. Rhodamine 123 accumulates in negatively charged compartments according to Nernst law.^37^ Accordingly, the mitochondrial membrane potential in milliVolts is given by the formula:

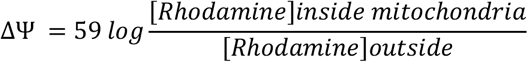

Accumulation of Rhodamine inside mitochondria causes quenching and decreases the fluorescence as mitochondrial membrane potential increases, experiments were as follow: Four identical additions of Rhodamine 123 solution were added to the mitochondrial suspension before initiation of respiration by succinate to reach a final concentration of 1 µM. This verified the linear relationship between Rhodamine concentration and fluorescent signal (Fluo) in the zero-1µM range and considered as a calibration. At the end of the experiment a “zero potential reference state” was obtained with addition of cyanide, its value (Fluo_zero_) was, as expected, quite close to that obtained with the final concentration of Rhodamine during calibration. The contribution of intramitochondrial (quenched) Rhodamine to fluorescence was neglected (null) and then the fluorescent signal was supposed to result from Rhodamine present outside mitochondria (Rho_ext_). Then the difference between measured fluorescence and that observed in the zero potential reference state indicated the quantity of Rhodamine internalized by mitochondria (Rho_mitoch_) and their ratio could be obtained with the formula:

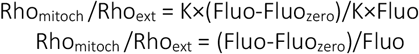

The external and intramitochondrial volumes had to be taken into account to obtain the ratio of concentrations according to the formula:

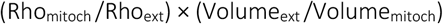

It was assumed that 1mg protein of the mitochondrial preparation corresponds to 0.5µL of mitochondrial internal volume (0.0005 mL/mg protein). Because working concentration of mitochondria are in the range of 0.5-1 mg/mL, mitochondrial volume is of negligible importance (≤1/1000) and the extramitochondrial volume is supposed equal to the volume of the experiment: 2mL of the O2K chamber. The ratio of these volumes is therefore given by the formula below, where “mg” is the quantity of mitochondrial proteins in the experiment.

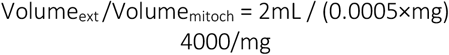

Hence the ratio between intramitochondrial and external Rhodamine concentrations could be expressed from fluorescence readings by the following expression:

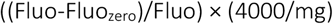

To be introduced in the Nernst equation above.

The fluorescence signal (a decrease) is a direct consequence of Rhodamine uptake by mitochondria that takes place according to Nernst’s law. Accordingly, a 180mV potential means a thousand times higher concentration of Rhodamine inside mitochondria. Starting from this maximal value of 180 mV a decrease in potential to 120 mV is expected to cause a division by ten of signal, a same factor is applied for decrease from 120 to 60 mV and again from 60 to 0mV. Then evaluation of membrane potential in the 60 to 120mV range would rely on variations within 10% of the maximal signal and from 0-60 mV within 1%. Moreover, measurement of the concentration dependence of Rhodamine fluorescence in the 0.5-1000µM range (not shown) indicated that quenching starts with concentrations above 10µM (60mV) and is complete (fluorescent signal becomes independent from concentration) above 60µM (105mV). This would made distinction between 60mV or zero impossible and assumption of a null fluorescence for intramitochondrial rhodamine is expected to result in inexact values below 105 mV.

### Rate of mitochondrial ATP production and calculation of ATP/O ratio

ATP/O appreciates the efficiency with which energy from the oxidation of substrates is converted into energy for ATP synthesis. Its determination requires the synchronous measurement of OCR and ATP production, the ratio between the latter and the former yields then the ATP/O value. Mitochondrial respiration was settled as above but after closure of chambers 1µM of P1, P5-Di(adenosine-5’) pentaphosphate (Ap5A) was added to inhibit any ATPase activity. In one chamber we measured OCR and take samples for ATP assay and in the other we measured membrane potential. 1 µM of Rho 123 was therefore added in this second chamber before initiation of respiration in both chambers with succinate (7.5mM). After the period for equilibration, TZ or RM were added to induce the required decrease in ΔΨ as judged from fluorescence measurement in the “rhodamine chamber”. Control, experiments included an equal volume of vehicle (DMSO). After initiation of ATP synthesis (Addition of ATP 5mM), 15 μL samples were taken 5 then 3 times in the interval of 30 and 60 seconds respectively. Two others samplings took place after addition of oligomycin with an interval of 2 min to estimate a possible ATP hydrolysis rate to be taken into account if present. We measured ATP concentration with the “ATP bioluminescence assay kit HS II” (Roche) according to the manufacturer instructions. Increase in ATP concentration in successive samples before oligomycin addition gave the ATP production rate expressed as nanomoles ATP produced per minute and milligram protein. ATP/O ratio was therefore calculated by dividing the ATP production rate by the OCR observed during the sampling phase.

### Statistical analysis

Statistical analyses were performed using Prism 9.0 software (GraphPad Software). Data in percentage are expressed as the percentage of corresponding control values. Mean values ± SEM/SD result from data points obtained in at least two independent experiments. Comparisons among groups were performed by one-way ANOVA followed by the Dunnett’s test. When all pairwise comparisons were carried out, one-way ANOVA was followed by the post hoc Tukey test. p values of <0.05 were considered significant.

